# Bird colours in urban mosaics: a study of two passerines

**DOI:** 10.1101/2025.07.15.664885

**Authors:** Katarzyna Janas, Marion Chatelain, Michela Corsini, Arnaud Da Silva, Łukasz Wardecki, Justyna Szulc, Marta Szulkin

**Affiliations:** Museum and Institute of Zoology, Polish Academy of Sciences, Warsaw, Poland; Institute of Environmental Sciences, Jagiellonian University, Poland; Department of Zoology, University of Innsbruck, Innsbruck, Austria; Institute for Alpine Environment, Eurac Research, Bolzano, Italy; Ecosphère, Mérignac, France; Polish Society for the Protection of Birds (OTOP); Institute of Evolutionary Biology, Faculty of Biology, Biological & Chemical Research Centre, University of Warsaw, Warsaw, Poland

**Keywords:** urban-dullness phenomenon, carotenoid-based colouration, phenotypic variation, structural colouration, melanin-based colouration, great tit, blue tit, tree cover, ISA

## Abstract

1. Urban environments are characterized by markedly altered conditions, which can exert a negative impact on the organisms that inhabit them. Among phenotypic traits most sensitive to urbanization is the colourful plumage of birds.
2. One of the best studied examples might be the urban dullness phenomenon, described for carotenoid-based traits, referring to colours being subdued in urban areas. In contrast, melanin-based and structural colouration are still understudied in the context of urbanization. Moreover, much research focused on changes in mean trait expression, while the effect on phenotypic variation, vital from an eco-evolutionary perspective, was rarely studied in this context.
3. Here, we examined urbanization-driven differences in expression and phenotypic variation of carotenoid-based, melanin-based and structural colours of two urban adapters, the great tit (*Parus major*) and the blue tit (*Cyanistes caeruleus*). Importantly, birds were sampled across multiple urban and suburban habitats, and replicated in eight Polish cities, located in the under-studied Central and Eastern European region.
4. We found a consistent decrease in mean carotenoid chroma of great tit breast plumage in more heavily urbanized habitats. This trait was also characterized by higher urban phenotypic variation, which could stem from greater environmental heterogeneity in cities. In blue tits, we observed reduced brightness of breast feathers in urban city centres, and an increased brightness of blue tail feathers in more urbanized habitats.
5. Our study sheds light on the complex pattern of colour trait sensitivity to urbanization and emphasizes the need for examining a wider range of species to gain greater insight into the eco-evolutionary processes acting in urban ecosystems.

## 1. Introduction

Urbanization profoundly transforms natural habitats, altering ecological conditions through increased pollution, habitat fragmentation, and changes in food availability, temperature, and light regimes (Grimm et al., 2008). These environmental changes pose physiological and ecological challenges to wildlife, even for species that appear well adapted to urban environments (Corsini et al., 2021, 2022; Corsini & Szulkin, 2025; Hahs et al., 2023). Among traits that are influenced by these urban stressors, animal colouration— particularly in birds—has received increasing attention. Plumage colouration plays crucial roles in thermoregulation, camouflage, species recognition, and sexual signalling (Hill, 2006b). It is shaped by a variety of mechanisms, including pigmentation (e.g., carotenoids and melanin), structural colouration, or a combination of both (McGraw, 2006a, 2006b; Prum, 2006). These mechanisms are differently sensitive to environmental variation, especially to changes in diet quality, stress levels, and pollutant exposure— factors that are often altered in urban settings (Hill, 2006a; Janas et al., 2024a). Accordingly, urban stressors can disrupt colour expression, as shown across multiple taxa, and particularly in birds (reviewed in Lifshitz and St Clair, 2016 and Leveau, 2021). Understanding how urbanization affects plumage coloration can provide insights into the ecological and evolutionary responses of animals to urban environments.

Undoubtedly, carotenoid-based colouration is the most extensively studied and most sensitive class of ornament to anthropogenic factors (Lifshitz and St Clair, 2016; Janas et al., 2024a). Carotenoid pigments cannot be synthesized by birds, thus their deposition in integumentary tissues is exclusively related to dietary intake (Hill, 1992). This links environmental quality, specifically the availability of carotenoid-rich food sources, to colour expression, typically measured as carotenoid chroma. In polluted or urbanized areas, the concentration of carotenoids might be lower in both primary carotenoid producers e.g., trees (Isaksson, 2009) and in leaf-eating caterpillars (Isaksson and Andersson 2007), which are important prey for many insectivorous passerines. This is thought to be the main reason for decreased chromaticity of carotenoid-based plumage of urban birds — known as ‘*urban dullness*’, most comprehensively documented in a recent multi-analytical study of Salmón et al., (2023). Moreover, the expression of carotenoid-based signals may also be mediated by oxidative stress, increased in urban environments (Giraudeau et al., 2015; Hutton and McGraw, 2016).

Despite the relative abundance of studies on the urban dullness phenomenon, these have focused on a restricted number of species, with the vast majority concentrating on the great tit (*Parus major*) (e.g., Grunst et al., 2020; Hõrak et al., 2000; Isaksson et al., 2005) and the house finch (*Haemorhous mexicanus*) (e.g., Giraudeau et al., 2015; Hasegawa et al., 2014; Hutton and McGraw, 2016; Sykes et al., 2021). Rare examples of research on other species include a study on the invasive common mynas (*Acridotheres tristis*) in replicated cities in Australia (Peneaux, Grainger, et al., 2021) and a study on the Eurasian kestrels (*Falco tinnunculus*) in Vienna (Sumasgutner et al., 2018). Crucially, no data from Central-Eastern European cities exist to date (but see Janas et al., 2024b for a natural cavity-nestbox comparison). Moreover, a recent study on northern cardinals (*Cardinalis cardinalis*) reported a reversed pattern, with more saturated red colouration in urban birds, due to higher access to invasive honeysuckle (Lonicera spp.) shrubs (Baldassarre et al. 2023). Therefore, to better understand the prevalence of *‘urban dullness’*, it is essential to conduct studies that include interspecific comparisons and replicated sampling designs, across multiple urban gradients.

Beside the chromatic component, affected by urban dullness phenomenon, an important dimension of carotenoid-based colouration is its achromatic component. It is most often quantified as brightness, and is thought to be determined by structure, i.e., density of barbs, variation in their keratin/air matrix or quality of keratin itself (Jacot et al., 2010; Shawkey & Hill, 2005). These properties, in turn, may depend on the quality of bird diet during moult (Mahoney et al., 2022). However, although the diet of urban birds differs from that of their rural/forest counterparts (Chatelain et al., 2025), this was not reflected in altered brightness of carotenoid-based feathers in most studied populations (Biard et al., 2017; Salmón et al., 2023; but see: Sandmeyer et al., 2025).

In the case of melanin-based colouration, a well-known phenomenon is urban melanism, which refers to the increased frequency of melanic morphs or patterns in more polluted habitats (Leveau, 2021; Lifshitz & St Clair, 2016). The most notable instance, described as industrial melanism, was documented in the Peppered moth (*Biston betularia*) during the Industrial Revolution in England (Cook et al., 2012). In birds, most evidence comes from studies on feral pigeons (*Columba livia*) with melanic morphs occurring more frequently in urban habitats (e.g., Jacquin et al., 2013; Obukhova, 2007). Apart from the modified frequency of dark morphs, anthropogenic factors like metal pollution or urban-related oxidative stress were also proposed to modify the intensity of melanin production and deposition, affecting feather brightness or ornament size (Galván & Alonso-Alvarez, 2009; Hutton & McGraw, 2016; McGraw, 2003). It was also hypothesized that melanized, dead tissues such as feathers can accumulate excess heavy metals, thereby protecting birds from their toxic effects (Chatelain et al., 2016; McGraw, 2003). However, despite numerous empirical attempts to demonstrate the impact of pollutants on the intensity of melanin-based traits, the results have been inconsistent. A recent meta-analysis summarizing research on the effects of heavy metals, persistent organic pollutants, and urbanization, revealed no significant impact on avian melanin-based traits (Janas et al., 2024a).

Finally, structural colouration (both, iridescent and non-iridescent types), has rarely been studied in the context of urbanization (unlike pigment-based ornaments). Chatelain et al., (2017) demonstrated that the feathers of feral pigeons exposed to higher levels of lead (both naturally in urban environments and in an experimental setting) had lower iridescent feather brightness. In the case of non-iridescent structural colouration, Tringali and Bowman (2015) compared UV-blue colouration of Florida Scrub-Jays (*Aphelocoma coerulescens*) from suburban and wildland populations. They reported that suburban birds developed more UV-reflective feathers, possibly due to greater food abundance, and that after immigrating to wildland sites, they outcompeted local birds. Despite this, they had lower reproductive success, which suggests that above-average availability of low-quality food in suburban sites provides a mismatch between trait expression and its signalling value (here: nutritional condition).

Beside the impact of urban-related environmental factors acting during the moulting period, which might affect the development and quality of produced feathers, their colour (as perceived by conspecifics) might be affected by air pollution adhering to the feather surface and altering their reflectance. Such indirect negative impact of urban-related airborne particles on the expression of structural colouration has been demonstrated in experimental studies on iridescent colouration of European starling (*Sturnus vulgaris*) (Griggio et al., 2011; Surmacki et al., 2023). Long-term exposure of starling feather samples to air-pollution resulted in significant reduction of reflectance, especially in the UV region (Griggio et al., 2011). Corresponding results were found in eastern bluebirds (*Siala sialis*), whose feathers, when exposed to the urban environment, had decreased values of UV and blue chroma (Surmacki et al., 2023). Given those findings, the lack of studies on the impact of urbanization on plumage ornaments with structural colouration, particularly in cities of Central and Eastern Europe where pollution levels often exceed EU ranges (European Environment Agency, 2019) represents a significant caveat.

Although multiple studies examined the impact of urban-related factors on avian plumage colour, reported values were usually centred around the detection of differences in phenotypic mean, and often by only comparing urban versus rural/forest populations. However, it is now established that urbanization can also influence phenotypic variation, potentially leading to important eco-evolutionary consequences in terms of selection and response to selection (Thompson et al., 2022). In urban populations, phenotypic variation might be influenced by several mechanisms, including relaxed selection and higher environmental heterogeneity within urban environments. Thus, a comprehensive analysis of 13 European populations of great tits and blue tits thus revealed a decrease in mean morphological traits and an increase in variance in urban populations (Thompson et al., 2022). However, to our knowledge, no studies have yet explored this issue in relation to avian colour traits.

In this study, we examined urbanization-driven differences in carotenoid-based, melanin-based and structural colour expression of two urban adapters, the great tit (*Parus major*) and the blue tit (*Cyanistes caeruleus*). To account for the spatial heterogeneity of urban landscapes (Szulkin et al., 2020), we sampled birds across six habitat types in urban – forest gradients, replicated in eight Polish cities. Moreover, to ensure comparability of urbanization levels with different studies (Leveau, 2021), we characterized urbanization intensity using percentage of impervious surface cover (ISA) and tree cover.

Specifically, we addressed two research questions:

1. How does urbanization affect plumage ornaments with different colour producing mechanisms?
2. Does urbanization modify the phenotypic variation of colour traits?

The inclusion of two study species, collected concurrently at the same sampling sites, allowed to examine a wider palette of colour traits and enabled a comparison of their sensitivity to urbanization in this aspect. Based on earlier studies (Salmón et al., 2023), we predicted to observe the urban dullness phenomenon in the carotenoid-based breast colouration of both great tits and blue tits. Consequently, we expected to find lower carotenoid chroma in more urbanized habitats, with no corresponding effect on feather brightness.

Moreover, considering the high heterogeneity of urban environments (Thompson et al., 2022) and the established link between carotenoid-based colouration and environmental pigment availability, we also predicted breast colouration to be more variable in more urbanized sites. Considering the contrasting results of previous studies inferring variation in melanin-based traits, and the scarcity of research on structural colouration in the context of urbanization, we bring forward new data on urban colouration from an understudied geographical location with specific urban historical and political legacy (Carlen et al. 2025).

## 2. Methods

### 2.1 Study species

Great tits and blue tits are closely related passerine species, abundant in forests and urban habitats. Although they occupy a similar ecological niche, there are some differences in their foraging strategy, reproductive ecology or sensitivity to urban conditions (Chatelain et al., 2025; Corsini et al., 2021; Milligan et al., 2017; Sudyka et al., 2022). Both species express conspicuous plumage patterns with carotenoid-based colouration on breast and belly. The great tit has melanin-based colouration with structural glow on cap, and white cheeks, while the blue tit expresses non-iridescent structural UV-blue colouration on crown, wing flight feathers and coverts and tail. Moreover, in the great tit, breast tie is known to be a signal of social dominance (e.g., Järvi and Bakken, 1984; Quesada and Senar, 2007), while the blue tit has only a narrow breast stripe, which is not known to play any signalling function, and was not analysed in this study.

### 2.2 Study sites

Great tits and blue tits were caught in 2018, in 44 study sites across urban-forest gradients in 8 Polish cities: Białystok, Katowice, Łódź, Toruń, Lublin, Poznań, Warszawa and Wrocław (Figure 1.A). For each city, sampling sites were chosen to represent each of the following urban habitats: city centres, residential areas, urban parks and suburban forests. Sampling locations for each type of habitat was triplicated in Warsaw – the capital city of Poland and largest city in the dataset. Additionally, in 4 cities, we established four sampling sites that were categorized as river corridors, with sampling sites in Warszawa and Toruń (Vistula river), Poznań (Warta river) and Wrocław (Odra river).

**Figure 1.**
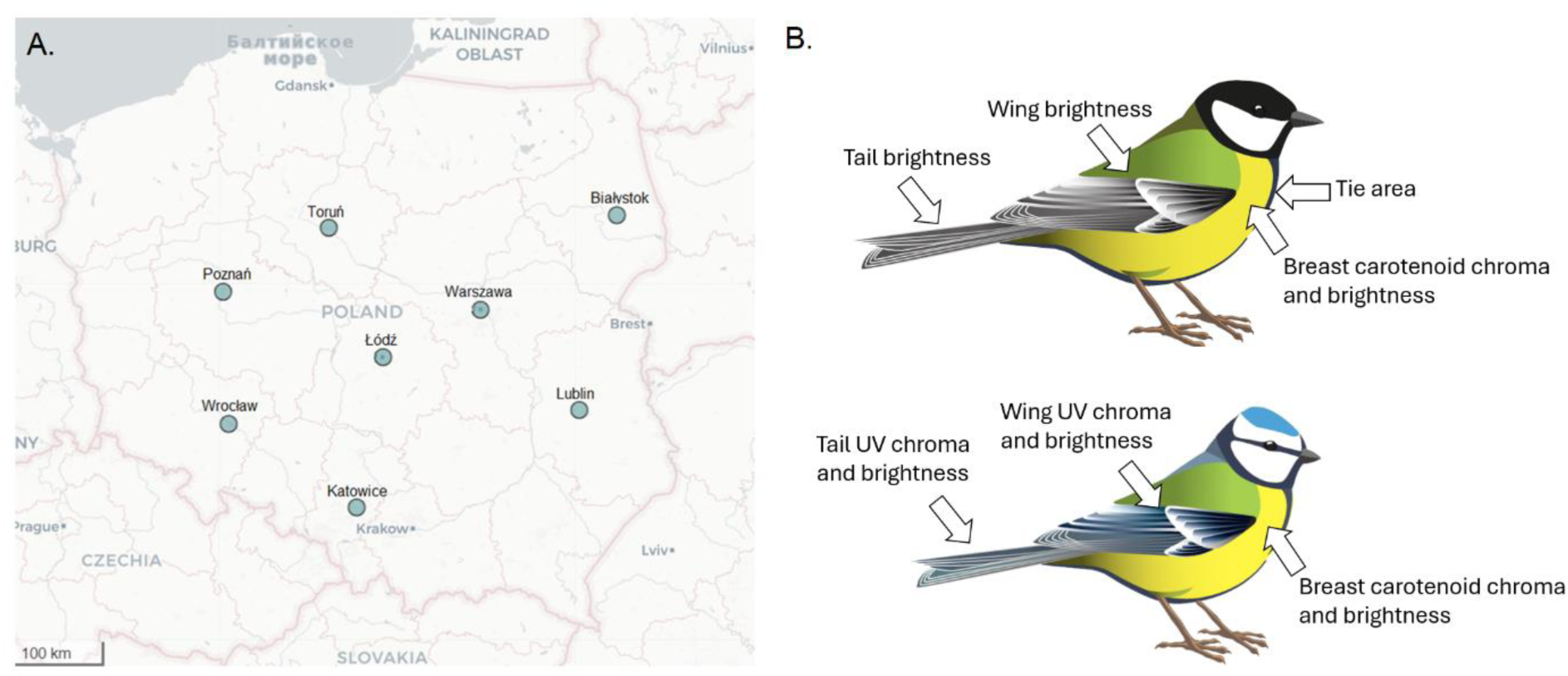
A. Map of cities where birds from a mosaic of habitats were sampled. In each city, birds were sampled in 4 to 13 sampling sites. **B.** Feather sampling location and type of colour trait analysed.

Following Chatelain et al. 2021, we further used two metrics characterizing the magnitude of urbanization for each sampling site: the percentage of impervious surfaces (ISA – impervious surface area) and tree cover within a radius of 100 meters from the sampling site. Averaged urbanization metrics for each habitat type are presented in Table 1.

**Table 1.** Summary statistics for the number of (i) study sites across 8 cities, (ii) sampled individuals and (iii) urbanization metrics for each habitat type (mean ± SE).

| Habitat type | Sites | Blue tits | Great tits | ISA (%) | Tree cover (%) |
| --- | --- | --- | --- | --- | --- |
| City centre | 10 | 15 | 37 | 52.09 ($\pm 1.57$) | 15.45 ($\pm 1.76$) |
| Residential area | 10 | 25 | 46 | 27.98 ($\pm 1.14$) | 16.88 ($\pm 1.87$) |
| River corridor | 4 | 13 | 13 | 6.82 ($\pm 2.0$) | 53.37 ($\pm 4.29$) |
| Urban park | 10 | 38 | 72 | 2.37 ($\pm 0.44$) | 50.98 ($\pm 2.22$) |
| Forest | 10 | 17 | 33 | 0.09 ($\pm 0.03$) | 72.77 ($\pm 1.89$) |

### 2.3 Bird sampling

The sampling took place between March 6th and April 11th, 2018. Birds were caught using mist-nets and blue tit and great tit call playbacks through loudspeakers. Each bird was ringed with individually numbered rings (Polish Ringing Centre, Ornithological Station, MiIZ PAS, Gdańsk, Poland). Birds were sexed based on brood patch presence and plumage features (the latter in great tits only) and aged (1^st^ year breeders or older) based on the moult limit between primary and greater coverts (Demongin 2016). Mass was measured with a digital scale to the nearest 0.1g. The length of the right tarsus was measured three times with a calliper, and the mean value was used for further analysis. Residuals from a regression of body mass on tarsus length were used as an index of current body condition (hereafter: condition) (Jakob et al., 1996). Breast feathers (approximately eight feathers), the second left tail feather, and the innermost secondary feather of the left wing were collected. Finally, great tit males were photographed with a digital camera (Nikon D3200 with AF-S Nikkor 18-55 mm lens) for tie area measurement (Nicolaus et al., 2016).

In total, we collected feather samples from 309 birds, 201 great tits (200 and 198 of wing and tail feather samples, respectively) and 108 blue tits (Table 1). Birds were caught at the early stage of the breeding season, which could have contributed to the significant male bias in sample size (females constituted 36,1% and 36,14% of caught individuals, respectively in great tits and blue tits).

### 2.4 Feather colouration analysis

Feather samples (Figure 1.B) were preserved in parchment envelopes and shielded from light. Shortly prior to the measurements, breast feathers were stuck on black paper in tiled stacks using double-adhesive transparent tape (ca. 8 feathers in a pile). Such samples were labelled with the bird’s ring number only, so that the measurements were blind to bird origin or sex. Wing and tail feathers were measured against black background paper, with measurements carried out on a fragment of the inner vane, at a height of 5 - 15 mm from the top of the feather. Reflectance was measured with the spectrophotometer Maya 2000 Pro with a PX-2 pulsed xenon light source (Ocean Optics, USA) and fiber-optic cable 6 × 400 µm with custom-made 2-mm collar. Measurements were repeated five times for tail and wing feathers, and ten times for breast and crown feathers, with a reference scan of white standard (Labsphere) taken every 10 minutes.

The spectra were processed, smoothed and averaged within each sample in the package *pavo2.0* (v. 2.9.0, Maia et al., 2019). For breast feathers of both species, we calculated mean brightness and carotenoid chroma ([R700 − R450]/R700). For structurally coloured crown, wing and tail feathers of the blue tit, we calculated mean brightness and UV chroma (R300−400/R300−700). Only mean brightness was calculated from melanic great tit wing and tail feathers (Figure 1.B). Within-sample repeatability of measurements was calculated in the package R-*rptR* (v. 0.9.22, Stoffel et al., 2017) and is presented in Table S1, which varied between 0.21 and 0.76 depending on the colour metric. Reasons for low repeatability values of wing and tail feather brightness stems from the fact that there is (i) uneven brightness in those feathers and (ii) that the probes were moved between measurements. Correlations between colour metrics within analysed patches are given in Figure S1.

The extent of black breast tie area were analysed with Gimp 2 version 2.8.14 (The GIMP Development Team, 2024) using photographs of 139 male great tits. Photographs were standardized using a black and a white reference on the picture. The tie area was measured using a slightly modified protocol from Nicolaus et al., (2016). It was calculated as the black surface comprising a 4-cm-long rectangle starting at the lowest point of yellow on the chest and as broad as the width of the belly.

### 2.5 Statistical analysis

All statistical analysis and graphs were generated in the R environment v. 4.4.1 (R Core Team, 2024).

We examined species-specific differences in carotenoid-based breast plumage colouration (i.e., mean brightness and carotenoid chroma) using linear models with colour metrics as a response and species, sex, and age (first-year breeders or older) as categorical predictors. Both carotenoid chroma (F1;301 = 50.06, p < 0.001) and brightness (F1;301 = 210.31, p < 0.001) was significantly higher in the great tit (Figure S2). Therefore, we decided to analyse the impact of urbanization on plumage colour traits separately for great tits and blue tits.

To analyse great tit and blue tit colour traits expression in relation to different urban habitat types, we tested a set of linear mixed effects models, fitted using *lmer* function in the *lmerTest* package (v. 3.1-3, Kuznetsova et al., 2017). The models included Z-score scaled colour metrics as response variables and habitat type (city centre, residential area, urban park, river corridor and forest), sex, age (birds in their second year or older) and condition as explanatory variables. The city (location) was introduced as a random intercept, to account for the variability resulting from local urban characteristics. To account for the possibility of differential response to urbanization between sexes and age classes, in each model we initially tested the interactions between habitat type and sex, age, and interaction between sex and age, which were sequentially removed when non-significant.

The models’ assumptions were tested using the package *DHARMa* (v. 0.4.7, Hartig, 2024). We observed significant quantile deviations in several models; appropriate transformations were consequently applied (great tit breast carotenoid chroma was squared, great tit tie area and blue tit breast carotenoid chroma were log-transformed). Conditional and marginal R^2^ were calculated using the package *performance* (v. 0.12.4, Lüdecke et al. 2021). In models with significant differences between habitat types, *post-hoc* least square mean pairwise comparisons were performed using *lsmeans* package (v. 2.30-0, Lenth, 2016). Moreover, to examine differences in phenotypic variation of colour traits between different habitat types, we performed the asymptotic test for equality of coefficients of variation (Feltz and Miller, 1996) using the package *cvequality* (Marwick and Krishnamoorthy, 2019).

A similar set of models was constructed to examine the relationship between colour traits and quantitative proxies of urbanization (i.e., tree cover and ISA). Given that tree cover and ISA were highly, negatively correlated (r = -0.70, df = 332, p < 0.001), their impact was analysed in separate models. The models were analogous as described above but included tree cover / ISA, instead of habitat type, in interaction with sex, and age. We detected quantile deviations in several models, thus appropriate transformations were applied in the response variables (see Table 3 and S3).

## 3. Results

### 3.1. Mean expression of colour traits in relation to habitat type

In great tits, we found significant differences in carotenoid-based breast plumage between different habitat types (Figure 2.A, Table S2). Specifically, birds from forest habitats had higher *carotenoid chroma* than those from city centres (Estimate ± SE; 0.69 ± 0.23, t = 3.00, p = 0.003), residential areas (Estimate ± SE; 0.49 ± 0.22, t = 2.23, p = 0.027) and urban parks (Estimate ± SE; 0.51 ± 0.20, t = 2.48, p = 0.014). While great tit breast feather *brightness* was significantly higher in males than in females, it did not significantly differ between habitats (Table S2). Similarly, we observed no significant differences in wing and tail *brightness* (which shows an inverse relationship with melanin concentration) between habitat types (Table S2). The breast tie area, analysed exclusively in males, was higher in older individuals and positively associated with condition but did not differ between habitat types (Table S2).

**Figure 2.**
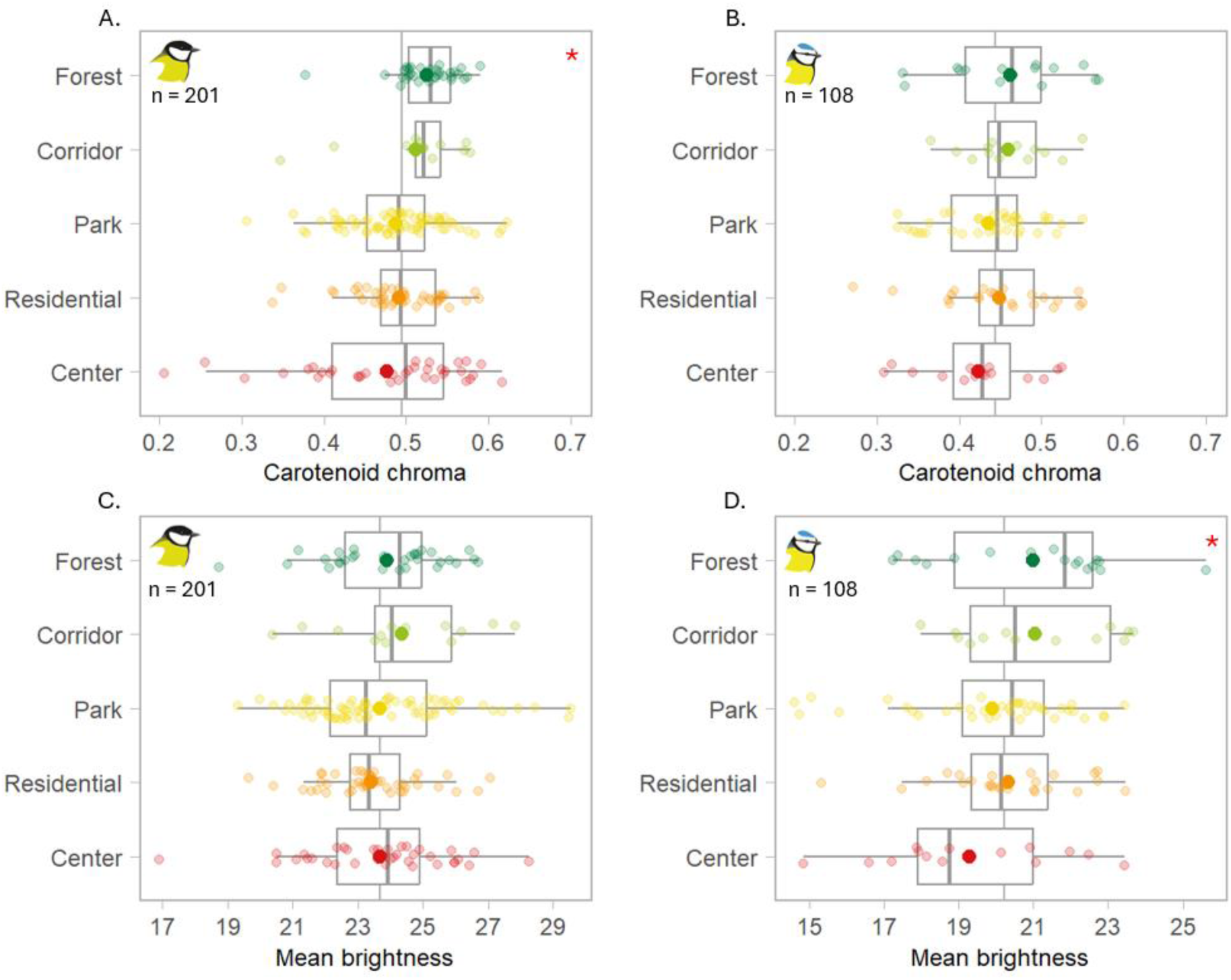
Boxplots for great tit and blue tit breast feather carotenoid chroma (A, B) and brightness (C, D) in relation to habitat type. Horizontal bars indicate data median, bigger dots indicate mean, box edges denote 25%–75% quartiles, whiskers indicate 1.5 IQR. Plotted on raw data. Red asterisk in the top right corner indicates significant differences between habitat types (Table S2).

In blue tits, breast *carotenoid chroma* plumage did not vary between different habitat types (Figure 2.B, Table S2). However, breast *brightness* in birds from forest habitats and river corridors was higher than in birds from city centres (respectively, Estimate ± SE; 0.77 ± 0.34, t = 2.27, p = 0.025 and 0.82 ± 0.37, t = 2.22, p = 0.029) and in males (Table S2). There were also differences in tail feather *brightness* between habitat types (Table S2, Figure 3), with forest birds expressing lower *brightness* than those from city centres and urban parks (respectively, centres: Estimate ± SE; -0.97 ± 0.32, t = -3.07, p = 0.003; urban parks: -0.62 ± 0.27, t = -2.22, p = 0.022) and between those from river corridors and city centres (Estimate ± SE; -0.71 ± 0.34, t = -2.07, p = 0.041). Tail *brightness* was also affected by the interaction between sex and age resulting from higher tail *brightness* in young females but the lack of sex difference in older individuals (Table S2, Figure S6). Sex- and age-specific interactions in wing *brightness* and *UV chroma* (respectively) were also observed (Table S2).

**Figure 3.**
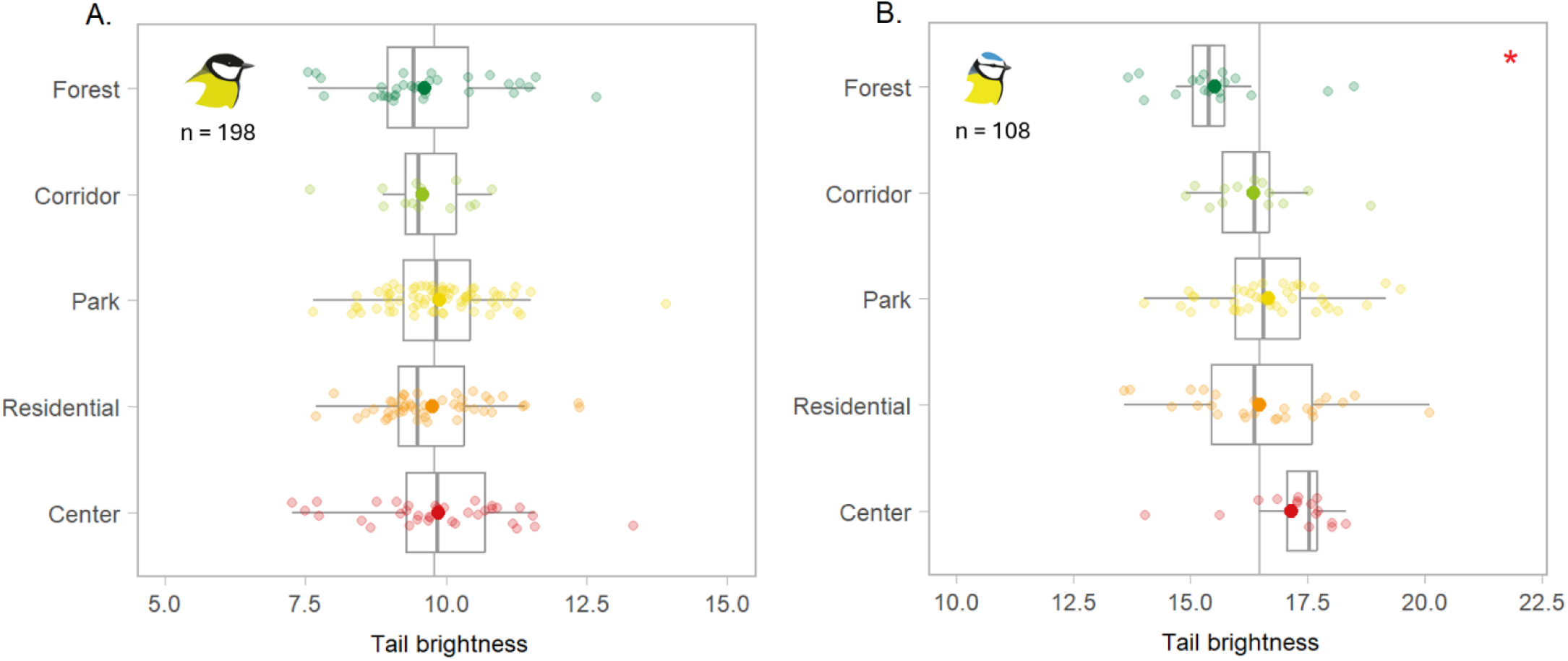
Boxplots of great tit (a) and blue tit (b) tail feather brightness in relation to habitat type. Horizontal bars indicate data median, bigger dots indicate mean, box edges denote 25%–75% quartiles, whiskers indicate 1.5 IQR. Plotted on raw data. Red asterisk in the top right corner indicates significant differences between habitat types in the models (Table 2).

### 3.2 Phenotypic variation of colour traits in relation to habitat type

Interestingly, there were significant differences in coefficients of variation (CVs) of great tits *carotenoid chroma* between different habitat types (test statistic = 36.73, k = 5, p < 0.001), with a clear tendency of increasing variation in more urbanised habitats (Figure 2). However, we found no differences in coefficients of variation between habitat type in any of the remaining great tit metrics and in any of the analysed blue tit metrics (Table S3).

### 3.3 Mean expression of colour traits in relation to tree cover and ISA

Given the similarity of model outcomes stemming from high negative correlation between the extent of tree cover and amount of impervious surface (ISA) across the sampling sites (Figure S3), the results of models examining the impact of ISA are presented in SI (Table S4). In great tits, there was a positive association between the amount of tree cover at the sampling site and breast feather plumage *carotenoid chroma* (Table S4, Figure 4.A). A similar relationship, with reversed direction and smaller effect size was observed for percentage of ISA (Estimate ± SE; 0.01 ± 0.003, t = -2.58, p = 0.011; Table S5). None of the other great tit colour metrics were related to tree cover and ISA (Table S4, Table S5).

**Figure 4.**
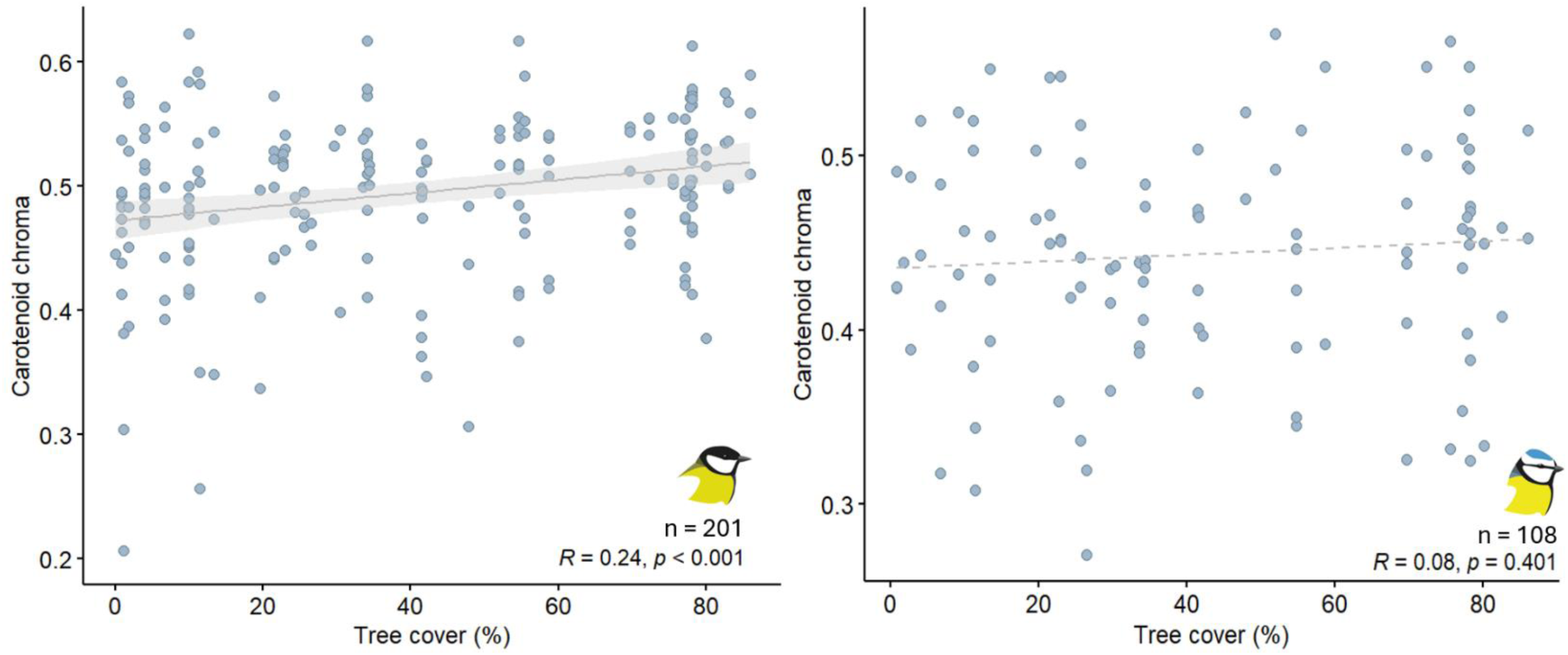
Scatterplots of raw data showing association between great tit (A) and blue tit (B) breast carotenoid chroma and tree cover (%).

**Figure 5.**
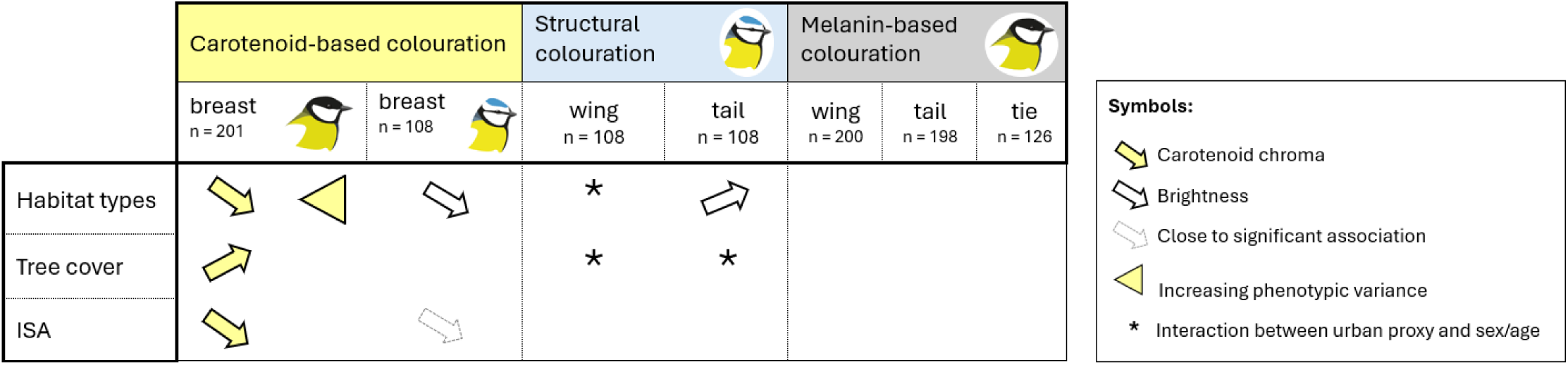
A graphic summary of main study outcomes.

In contrast to great tits, there was no association between tree cover or the amount of impervious surface with blue tit breast feather *carotenoid chroma* (Table S4; Table S5). However, there was a tendency towards a negative association between ISA and breast feather *brightness* (Table S5). Tail feather *brightness* was explained by the interaction between tree cover and sex (Estimate ± SE; -0.01 ± 0.01, t = -2.03, p = 0.045), with males expressing lower values (possibly caused by higher melanin deposition) in habitats with higher tree coverage (Table S4, Figure S4). Moreover, the association between tail UV chroma and tree cover was age-specific (Estimate ± SE; 0.01 ± 0.005, t = 2.15, p = 0.034), with higher differences between age classes in more forested habitats (Table S4, Figure S5.A). Similarly, there was an interaction between ISA and age in the model examining tail UV chroma, with increasing ISA leading to wanning differences between older and younger individuals (Table S5, Figure S5.B).

## 4. Discussion

Based on eight replicated urban gradients, we examined the impact of urbanization on colour traits in two model passerine species, the great tit and the blue tit. Our results strongly support previous findings on the prevalence of the ‘*urban dullness’* phenomenon in carotenoid-based colouration of great tit (Salmón et al., 2023). Thus, this replicated dataset of 8 urban mosaics confirms a reduction of breast feather carotenoid chroma in great tits. Moreover, we also report increased phenotypic variance of great tit breast carotenoid chroma in more urbanized habitats. However, in contrast to predictions, the analogous effect was not observed in the blue tit. Instead, breast feather brightness consistently decreased along the rural-urban gradient. We also examined the influence of urbanization on feather traits that are less often studied, including melanin-based colouration of the great tit and non-iridescent UV-blue structural colouration of the blue tit. Importantly, we observed a consistent increase in blue tit tail feathers brightness in more urbanized habitats. At the same time, and in contrast to earlier work (Senar et al., 2014), we did not find any evidence for an effect of urbanisation on great tit melanin-based breast tie, wing and tail feather colour traits.

### 4.1. Carotenoid-based colouration

In line with predictions, we found a negative impact of urbanization on the carotenoid chroma of breast plumage in the great tit (Figure 2.A). An analogous effect was not observed for feather brightness, interpreted as a measure of feather structural quality (Jacot et al., 2010; Shawkey & Hill, 2005), and is in line with earlier studies (Biard et al., 2017; Salmón et al., 2023; but see Sandmeyer et al. 2025, *preprint*). Carotenoid chroma depends on the amount of pigment deposited in the feather, thus, one of the important factors that shape its expression is environmental availability of carotenoid dietary sources. In great tits and blue tits, the main carotenoids are lutein and zeaxanthin, derived from invertebrates e.g., *Lepidoptera* caterpillars, which obtain carotenoids from the leaves of the trees such as oak (*Quercus robur*), birch (*Betula spp.*), and alder (*Alnus spp.*) on which they feed (Isaksson & Andersson, 2007; Partali et al., 1987). Thus, the availability of carotenoid-rich food in urbanized environments is limited by the extent of parks and urban forests. Accordingly, Chatelain et al., (2025) found that, in Innsbruck’s population, urban great tits consumed fewer arthropods (including Lepidoptera) during the breeding season compared to their rural counterparts. Although this pattern was no longer maintained in the following months, it is possible that the deficiencies experienced during reproduction have prolonged effects during the resource-demanding moult period. In our study, we found a positive relationship between tree cover and carotenoid chroma in great tits (Figure 4.A). The negative effects of limited availability of green spaces in cities may be further aggravated by heavy metal pollution, which can reduce both the carotenoid content and abundance of caterpillars (Isaksson and Andersson, 2007; Eeva et al., 2009; Isaksson, 2009), as well as interfere with pigment deposition (Peneaux, Hansbro, et al., 2021). Supporting this, Chatelain et al., (2021) found that in the same study sites as the present research, metal pollution was positively associated with the degree of urbanization — highlighting metal pollution as a strong candidate for explaining urban-driven variation in plumage coloration.

In addition to reduced mean carotenoid chroma in great tit, our study also reveals increased phenotypic variance of this trait in more urbanized habitats. To our knowledge, there are no earlier reports of colour trait variance in other studies of this type. Complementarily, an analogous pattern was described for morphological traits of great and blue tits from 13 pairs of urban-rural populations pairs across Europe (Thompson et al., 2022). The most plausible explanation for our findings is that high urban environmental heterogeneity, characterized by fragmented green areas interspersed with extensive impervious surface areas (ISA), induces higher variance in colour trait expression in urban birds. A mosaic of areas with increased local intensity of the ‘*urban heat island’* effect, may further exacerbate the issue of caterpillar’s availability (Visser and Gienapp, 2019). Moreover, despite the clear environment dependence of carotenoid-based colouration, Evans and Sheldon (2012) reported significant (though moderate in strength) heritability of yellow plumage chromaticity in great tits. They also showed that this heritability was not inflated by shared environmental factors, which suggests that foraging abilities may be heritable. This implies that the observed effects of urbanization on colouration may result from selection. Interestingly, significant heritability in the chromatic component of yellow colouration has also been described in blue tit males (with h^2^ between 0.13 and 0.25 in Corsica and Rouvière populations, respectively; Charmantier et al., 2017). However, in our study, we found no influence of urbanization on the variation in carotenoid-based chroma in blue tits.

In contrast to what was observed in the great tit, there was no significant effect of urbanization on the mean expression of breast plumage carotenoid chroma in blue tits. In both species, yellow colouration is based on the same dietary pigments, lutein and zeaxanthin, deposited in similar ratio (Partali et al., 1987). However, blue tit colouration is characterized by significantly lower carotenoid chroma, compared to great tits (Figure S2), which suggests that they require less pigment. It is therefore possible that environmental availability of carotenoid-rich dietary sources confined in more urbanized habitats (Isaksson & Andersson, 2007) is not as limiting a factor for colour trait expression as it is for great tits. Another, non-exclusive explanation, might be diet differences between the two species. In Innsbruck, while urban great tits shifted to a more opportunistic and granivorous diet (including anthropogenic food sources like sunflower seeds and peanuts), urban blue tits remained largely insectivorous from spring to autumn, thus potentially maintaining their carotenoid intake (Chatelain et al., 2025).

Interestingly, blue tits from city centres displayed lower brightness than birds from forests and river corridors (Figure 2.D). A similar, marginally non-significant tendency was found in female great tits in the long-term study from Montpellier (Sandmeyer et al. 2025, *preprint*). Given that brightness is thought to reflect the feather structural component i.e. microstructure at barb and barbule levels) and keratin quality (Shawkey and Hill, 2005; Jacot et al., 2010), the observed effect might be caused by the low nutritional condition of birds in the cities. Thus, McGlothlin et al., (2007) experimentally demonstrated that food composition, specifically its protein content, affects the size and brightness of white feather ornaments. Moreover, Chatelain et al., (2025) reported that blue tit in spring, urban individuals consumed less moths but more aphids and crab spiders, while from August to October, they ate fewer seed bugs but again more aphids, along with weevils and moths. These findings suggest that while blue tits remain largely insectivorous, the types of arthropods they consume—and consequently their protein intake—may vary along rural-urban gradients. In addition, lower brightness may also result from altered moult timing and duration in the cities (Hutton et al., 2021), as well as heightened exposure to oxidative stress and chemical pollution in cities (Isaksson, 2015).

The observed disparities in sensitivity to anthropogenic factors between the two studied species examined in this study highlight the fact that results pertaining to the *‘urban dullness’* phenomenon should not be extrapolated to even closely related species sharing the same habitats. Given the prevalence of carotenoid-based ornaments in birds and their importance in sexual selection (Delhey et al., 2023), the impact of urbanisation on feather colour traits warrants further investigation in a wider range of species.

### 4.2. Structural colouration

Our study represents one of the first attempts to examine the impact of urbanization on structural UV-blue colouration present in blue tit wing and tail feathers (as well as in the crown, not measured in this study). Two previous experimental studies demonstrated that long-term exposure of structurally coloured feathers to urban air pollution decreases their short-wavelength reflectance due to the absorption properties of airborne particles (Griggio et al., 2011; Surmacki et al., 2023). The design of these experiments, based on feather samples, was intended to eliminate the impact of other factors, including active feather preening. In our study, the most pronounced effect was an increased brightness of blue tit tail feathers in more urbanized sites (Figure 3.A). Given the direction of this effect, it can be ruled out that it was caused by the deposition of pollution on the feather surface and indicate that birds are capable of efficiently removing urban dust during preening. It appears that observed differences in brightness might arise rather due to the impact of urban-related factors on feather wear or bird physiology during moulting, which determines the quality of the emerging feather.

Similarly, as in the case of carotenoid-based colouration, a possible cause of the observed effects reported in this study (an increase in blue tit tail structural colouration brightness in urban birds) might be related to an increased abundance of anthropogenic food sources in urban habitats. A previous study with experimental food supplementation showed that male nestlings of eastern bluebirds (*Siala sialis*) raised by supplemented parents, developed blue primaries with higher brightness (Doyle and Siefferman, 2014). On the other hand, Florida Scrub-Jay (*Aphelocoma coerulescens*) from suburban sites with high availability of anthropogenic foods, developed feathers with higher UV chroma, but not brightness, than their wildland counterparts (Tringali & Bowman, 2015). Curiously, in our study, increased brightness in more urbanized habitats was observed only in tail but not in wing feathers. We presume that the investment of resources in the structural component of wing feathers is prioritized over remaining feathers, given their role in flight, and thus environmental effects might be less visible in this trait.

There was also an age-specific association between tree cover and tail UV chroma, with differences between age classes more pronounced in habitats with higher tree cover (Figure S5). An analogous interaction was present in the case of ISA, indicating that the primacy, in terms of ornament elaboration, of older individuals is fading with increasing urbanization. Thus, easier access to anthropogenic food sources in more urbanised habitats may eliminate the advantage that older, more experienced individuals have in competition for food in natural habitats. As suggested by Tringali and Bowman (2015) this may lead to a mismatch between signalled and actual bird quality, possibly resulting in maladaptive mate choices.

Importantly, pinpointing the proximal causes of variation in blue feathers is inevitably complex as their colour is determined by many components: In terms of microstructural parameters, brightness is related to melanin/keratin ratio, barb cortex thickness and spatial frequency of the spongy layer (Fan et al., 2019; Hegyi et al., 2024). At the level of macrostructure, it might be linked with the number of barbs or blue section length (Hegyi et al., 2018). Further urban-based longitudinal studies that would include the analysis of feather macro- and microstructure, are necessary to confirm observed effects and explain its causes.

### 4.3. Melanin-based colouration

In contrast to observed effects of urbanisation on carotenoid-based colouration, we found no support for the impact of urbanization on the expression of melanin-based traits. In the great tit, breast tie is considered to signal social dominance (Järvi and Bakken, 1984), individual quality (Norris, 1990) and personality traits such as aggressiveness, boldness and fast exploration (Quesada and Senar, 2007). Therefore, differences in tie area associated with the degree of habitat urbanization would suggest directional selection on personality traits correlated with this trait, potentially leading to local adaptation. In line with this expectation, breast tie width was shown to be lower in the city of Barcelona, compared to the birds from forest population (Senar et al., 2014). However, in our study, we found no differences in tie area between birds from different habitat types. Similarly, the was no effect of urbanisation on breast tie measured across 13 cities in the Czech Republic (Bauerová et al., 2017). Moreover, Grunst et al., (2020) also found that metal pollution and proximity to roads have no effect on great tit tie area (but see Dauwe and Eens (2008) for contrasting results). Furthermore, if melanin deposited in feathers serves a detoxication function by binding and storing metal ions, increased melanin deposition should be observed also in non-ornamental traits. Therefore, besides tie, we also examined the brightness of tail and wing feathers in the analyses, yet none of them were related with urbanization. Taken together, it suggests that great tit melanin-based traits appear to be unaffected by urban-related factors.

## 5. Conclusion

By observing the effect of replicated urban gradients, we demonstrate that the *‘urban dullness’* phenomenon is affecting urban great tits, impacting not only mean trait expression but also the phenotypic variance of carotenoid colouration, potentially having important eco-evolutionary implications. Moreover, our study revealed that great tit and blue tit colour traits respond to urban related factors in a species-specific manner. This result highlights that even closely related species, with overlapping ecological niches, can differ in feather colour sensitivity to anthropogenic stressors. Thus, any extrapolation of results on the impact of urbanization on colour-related phenotypic traits across species should be made with caution. We recommend that future studies on the impact of anthropogenic factors on avian colour traits should target a broader range of species and ornamental colours.

## Acknowledgements

We are very grateful to Marta Celej for her help in preparing documentation and application for ethical permits. We would like to thank Szymon Drobniak for providing access to the spectrophotometer. The study was funded with a Sonata Bis Grant (2014/14/E/NZ8/00386) from the Polish National Science Centre awarded to Marta Szulkin.

## Data availability statement

Raw data are available on the Figshare platform (https://figshare.com/s/26f43429d813bfeba925).

## Ethical Permits

Bird sampling was conducted based on the approval of the Local Ethical Committee nr I for Animal Experimentation in Warsaw (permission no. 220/2016).

## Author Contributions

Conceptualization: Marta Szulkin, Arnaud Da Silva, Marion Chatelain, Katarzyna Janas. Data curation: Katarzyna Janas, Marta Szulkin. Formal analysis: Katarzyna Janas. Funding acquisition: Marta Szulkin. Investigation: Katarzyna Janas, Justyna Szulc, Łukasz Wardecki, Michela Corsini, Marion Chatelain. Methodology: Katarzyna Janas, Michela Corsini, Marta Szulkin. Project administration: Marta Szulkin. Resources: Marta Szulkin. Software: Katarzyna Janas. Supervision: Marta Szulkin. Validation: Marta Szulkin. Visualization: Katarzyna Janas. Writing – original draft: Katarzyna Janas. Writing – review & editing: Marion Chatelain, Michela Corsini, Arnaud Da Silva, Łukasz Wardecki, Justyna Szulc, Marta Szulkin.

