## Supplement for "Bird colours in urban mosaics: a study of two passerines"

### Supplementary information

### Tables

**Table S1.** Repeatability colour metrics measurements within sample (10 measurements per sample). Brightness and carotenoid chroma of breast feathers were calculated for joint set of great tit and blue tit samples. R was calculated in the package "*rptR*" (Stoffel et al. 2017), SE and 95% CI were estimated with 1000 bootstraps.

| colour metric | n | R | SE | 95% CI | p |
|---|---|---|---|---|---|
| Breast brightness | 331 | 0.67 | 0.02 | 0.63 - 0.70 | <0.001 |
| Breast carotenoid chroma | 331 | 0.76 | 0.02 | 0.73 - 0.79 | <0.001 |
| GT wing brightness | 217 | 0.21 | 0.023 | 0.17 - 0.26 | <0.001 |
| BT wing brightness | 115 | 0.21 | 0.031 | 0.14 - 0.26 | <0.001 |
| BT wing UV chroma | 115 | 0.66 | 0.033 | 0.59 - 0.72 | <0.001 |
| GT tail brightness | 214 | 0.24 | 0.024 | 0.19 - 0.29 | <0.001 |
| BT tail brightness | 115 | 0.27 | 0.035 | 0.20 - 0.34 | <0.001 |
| BT tail UV chroma | 115 | 0.65 | 0.033 | 0.58 - 0.71 | <0.001 |

**Table S2**. Linear mixed models of t great tit and blue tit feather colour traits in relation to habitat type. The forest habitat is used as reference value for habitat type. Significant values are highlighted in bold.

| GREAT TIT | Estimate | SE | df | t value | p | |
|---|---|---|---|---|---|---|
| *Breast carotenoid chroma*^2 | | | | | | |
| Intercept | 0.17 | 0.23 | 62.43 | 0.756 | 0.4523 | |
| **Habitat type (Park)** | **-0.51** | **0.20** | **189.92** | **-2.482** | **0.01393** | * |
| Habitat type (Corridor) | -0.23 | 0.33 | 191.72 | -0.711 | 0.47797 | |
| **Habitat type (Residential)** | **-0.49** | **0.22** | **190.64** | **-2.226** | **0.02716** | * |
| **Habitat type (Centre)** | **-0.69** | **0.23** | **190.94** | **-2.998** | **0.00308** | ** |
| Sex (M) | 0.28 | 0.15 | 191.89 | 1.837 | 0.0678 | . |
| Age (older) | 0.08 | 0.15 | 191.52 | 0.515 | 0.60739 | |
| Condition | 0.12 | 0.07 | 191.21 | 1.710 | 0.08892 | . |
| Random effects: | | Variance | | | | |
| | Location | 0.06 | | Conditional $R^2$: 0.147 | | |
| | Residual | 0.90 | | Marginal $R^2$: 0.090 | | |
| *Breast brightness* | | | | | | |
| Intercept | -0.41 | 0.21 | 78.49 | -1.97 | 0.052 | . |
| Habitat type (Park) | -0.02 | 0.20 | 191.75 | -0.11 | 0.915 | |
| Habitat type (Corridor) | 0.18 | 0.32 | 190.89 | 0.55 | 0.581 | |
| Habitat type (Residential) | -0.12 | 0.22 | 191.95 | -0.54 | 0.593 | |
| Habitat type (Centre) | -0.11 | 0.23 | 191.92 | -0.46 | 0.646 | |
| **Sex (M)** | **0.64** | **0.15** | **179.82** | **4.32** | **<0.001** | *** |
| Age (older) | 0.05 | 0.15 | 159.50 | 0.36 | 0.717 | |
| Condition | 0.07 | 0.07 | 184.24 | 0.97 | 0.335 | |
| Random effects: | | Variance | | | | |
| | Location | 0.01 | | Conditional $R^2$: 0.123 | | |
| | Residual | 0.91 | | Marginal $R^2$: 0.117 | | |
| *Wing brightness* | | | | | | |
| Intercept | 0.23 | 0.22 | 59.97 | 1.02 | 0.310 | |
| Habitat type (Park) | -0.01 | 0.20 | 188.78 | -0.05 | 0.963 | |
| Habitat type (Corridor) | -0.01 | 0.32 | 190.77 | -0.04 | 0.968 | |
| Habitat type (Residential) | 0.14 | 0.22 | 189.63 | 0.66 | 0.509 | |
| Habitat type (Centre) | 0.38 | 0.23 | 189.97 | 1.67 | 0.097 | . |
| **Sex (M)** | **-0.54** | **0.15** | **190.72** | **-3.62** | **<0.001** | *** |
| Age (older) | -0.10 | 0.15 | 190.22 | -0.68 | 0.499 | |
| Condition | 0.01 | 0.07 | 189.85 | 0.11 | 0.916 | |
| Random effects: | | Variance | | | | |
| | Location | 0.05 | | Conditional $R^2$: 0.145 | | |
| | Residual | 0.87 | | Marginal $R^2$: 0.093 | | |
| *Tail brightness* | | | | | | |
| Intercept | 0.53 | 0.21 | 67.71 | 2.50 | 0.015 | * |

| | | | | | | |
|---|---|---|---|---|---|---|
| Habitat type (Park) | 0.09 | 0.20 | 187.54 | 0.45 | 0.651 | |
| Habitat type (Corridor) | 0.17 | 0.32 | 188.71 | 0.54 | 0.592 | |
| Habitat type (Residential) | -0.02 | 0.22 | 188.52 | -0.08 | 0.935 | |
| Habitat type (Centre) | 0.16 | 0.22 | 188.79 | 0.71 | 0.480 | |
| **Sex (M)** | **-0.70** | **0.15** | **185.51** | **-4.83** | **<0.001** | *** |
| **Age (older)** | **-0.29** | **0.15** | **177.33** | **-2.00** | **0.047** | * |
| Condition index | 0.12 | 0.07 | 184.36 | 1.72 | 0.087 | . |
| Random effects: | | Variance | | | | |
| | Location | 0.02 | | Conditional $R^2$: 0.180 | | |
| | Residual | 0.85 | | Marginal $R^2$: 0.157 | | |
| <u>*Log Tie area*</u> | | | | | | |
| Intercept | -0.02 | 0.22 | 49.85 | -0.07 | 0.944 | |
| Habitat type (Park) | -0.26 | 0.25 | 111.74 | -1.01 | 0.313 | |
| Habitat type (Corridor) | -0.21 | 0.38 | 117.99 | -0.54 | 0.587 | |
| Habitat type (Residential) | -0.18 | 0.28 | 114.58 | -0.63 | 0.528 | |
| Habitat type (Centre) | -0.31 | 0.28 | 118.00 | -1.09 | 0.277 | |
| Age (older) | 0.38 | 0.18 | 80.91 | 2.11 | 0.038 | * |
| Condition | 0.23 | 0.09 | 108.56 | 2.42 | 0.017 | * |
| Random effects: | | Variance | | | | |
| | Location | 0.001 | | Conditional $R^2$: 0.084 | | |
| | Residual | 0.953 | | Marginal $R^2$: 0.084 | | |

| **BLUE TIT** | **Estimate** | **SE** | **df** | **t value** | **p** | |
|---|---|---|---|---|---|---|
| *Log Breast carotenoid chroma* | | | | | | |
| Intercept | -0.17 | 0.32 | 50.04 | -0.54 | 0.591 | |
| Habitat type (Park) | -0.33 | 0.30 | 98.00 | -1.10 | 0.274 | |
| Habitat type (Corridor) | 0.14 | 0.37 | 81.17 | 0.37 | 0.711 | |
| Habitat type (Residential) | -0.17 | 0.32 | 97.95 | -0.52 | 0.603 | |
| Habitat type (Centre) | -0.44 | 0.36 | 94.79 | -1.21 | 0.228 | |
| Sex (M) | 0.31 | 0.21 | 94.85 | 1.48 | 0.141 | |
| Age (older) | 0.33 | 0.20 | 80.22 | 1.67 | 0.098 | . |
| Fat | -0.25 | 0.21 | 86.34 | -1.16 | 0.250 | |
| Random effects: | | Variance | | | | |
| | Location | 0.001 | | Conditional $R^2$: 0.086 | | |
| | Residual | 0.986 | | Marginal $R^2$: 0.085 | | |
| <u>*Breast brightness*</u> | | | | | | |
| Intercept | 0.03 | 0.33 | 69.78 | 0.10 | 0.923 | |
| Habitat type (Park) | -0.46 | 0.28 | 95.66 | -1.63 | 0.107 | |
| Habitat type (Corridor) | 0.05 | 0.36 | 98.00 | 0.14 | 0.891 | |
| Habitat type (Residential) | -0.32 | 0.31 | 95.52 | -1.05 | 0.299 | |
| **Habitat type (Centre)** | **-0.77** | **0.34** | **94.88** | **-2.27** | **0.025** | * |
| **Sex (M)** | **0.47** | **0.20** | **95.53** | **2.39** | **0.019** | * |
| Age (older) | -0.18 | 0.19 | 98.00 | -0.94 | 0.348 | |

| | | | | | | |
|---|---|---|---|---|---|---|
| Condition | 0.01 | 0.21 | 97.71 | 0.04 | 0.967 | |
| Random effects: | | Variance | | | | |
| | Location | 0.120 | | Conditional $R^2$: 0.234 | | |
| | Residual | 0.852 | | Marginal $R^2$: 0.126 | | |
| *Wing brightness* | | | | | | |
| Intercept | 0.40 | 0.38 | 78.27 | 1.07 | 0.290 | |
| Habitat type (Park) | -0.18 | 0.30 | 96.48 | -0.59 | 0.560 | |
| Habitat type (Corridor) | -0.25 | 0.39 | 92.70 | -0.64 | 0.523 | |
| Habitat type (Residential) | -0.13 | 0.33 | 95.99 | -0.41 | 0.686 | |
| Habitat type (Centre) | 0.03 | 0.36 | 94.47 | 0.07 | 0.945 | |
| Sex (M) | -0.46 | 0.30 | 96.28 | -1.53 | 0.130 | |
| Age (older) | -0.57 | 0.34 | 96.98 | -1.68 | 0.097 | . |
| Condition | 0.01 | 0.22 | 95.16 | 0.03 | 0.973 | |
| **Sex (M): Age (older)** | **1.04** | **0.42** | **94.54** | **2.46** | **0.016** | * |
| Random effects: | | Variance | | | | |
| | Location | 0.036 | | Conditional $R^2$: 0.098 | | |
| | Residual | 0.996 | | Marginal $R^2$: 0.065 | | |
| *Wing UV chroma* | | | | | | |
| Intercept | -1.19 | 0.47 | 91.68 | -2.54 | 0.013 | * |
| **Habitat type (Park)** | **1.10** | **0.49** | **91.32** | **2.24** | **0.028** | * |
| Habitat type (Corridor) | 0.91 | 0.58 | 89.56 | 1.58 | 0.119 | |
| Habitat type (Residential) | -0.27 | 0.51 | 88.36 | -0.53 | 0.601 | |
| Habitat type (Centre) | 0.40 | 0.54 | 90.17 | 0.74 | 0.464 | |
| Sex (M) | 0.62 | 0.49 | 89.24 | 1.27 | 0.209 | |
| **Age (older)** | **1.02** | **0.18** | **93.89** | **5.74** | **<0.001** | *** |
| Condition index | 0.22 | 0.19 | 93.81 | 1.16 | 0.248 | |
| Habitat type (Park): Sex (M) | -0.99 | 0.57 | 90.32 | -1.74 | 0.086 | . |
| Habitat type (Corridor): Sex (M) | -0.94 | 0.69 | 89.64 | -1.37 | 0.173 | |
| Habitat type (Residential): Sex (M) | 0.51 | 0.60 | 87.58 | 0.86 | 0.393 | |
| Habitat type (Centre): Sex (M) | -0.23 | 0.66 | 89.75 | -0.36 | 0.723 | |
| Random effects: | | Variance | | | | |
| | Location | 0.081 | | Conditional $R^2$: 0.367 | | |
| | Residual | 0.684 | | Marginal $R^2$: 0.292 | | |
| *Tail brightness* | | | | | | |
| Intercept | 0.24 | 0.33 | 78.94 | 0.73 | 0.465 | |
| **Habitat type (Park)** | **0.62** | **0.27** | **95.60** | **2.33** | **0.022** | * |
| Habitat type (Corridor) | 0.26 | 0.34 | 95.61 | 0.77 | 0.442 | |
| Habitat type (Residential) | 0.56 | 0.28 | 95.07 | 1.97 | 0.052 | . |
| **Habitat type (Centre)** | **0.97** | **0.32** | **93.86** | **3.07** | **0.003** | ** |
| Sex (M) | -0.61 | 0.26 | 94.97 | -2.30 | 0.024 | * |

| | | | | | | |
|---|---|---|---|---|---|---|
| **Age (older)** | **-1.22** | **0.30** | **96.80** | **-4.08** | **0.000** | *** |
| Condition | -0.04 | 0.19 | 96.75 | -0.18 | 0.856 | |
| Sex (M): Age (older) | 0.88 | 0.37 | 93.76 | 2.40 | 0.018 | * |
| Random effects: | | Variance | | | | |
| | Location | 0.053 | | Conditional $R^2$: 0.304 | | |
| | Residual | 0.747 | | Marginal $R^2$: 0.254 | | |
| _Tail UV chroma_ | | | | | | |
| Intercept | -1.25 | 0.25 | 71.15 | -4.99 | <0.001 | *** |
| Habitat type (Park) | 0.28 | 0.22 | 97.03 | 1.25 | 0.214 | |
| Habitat type (Corridor) | -0.17 | 0.28 | 94.57 | -0.60 | 0.549 | |
| Habitat type (Residential) | 0.37 | 0.24 | 96.78 | 1.54 | 0.128 | |
| Habitat type (Centre) | 0.18 | 0.27 | 94.97 | 0.66 | 0.511 | |
| **Sex (M)** | **0.71** | **0.16** | **97.38** | **4.56** | **<0.001** | *** |
| **Age (older)** | **1.22** | **0.15** | **94.01** | **8.14** | **<0.001** | *** |
| Condition index | -0.05 | 0.16 | 96.85 | -0.32 | 0.754 | |
| Random effects: | | Variance | | | | |
| | Location | 0.022 | | Conditional $R^2$: 0.489 | | |
| | Residual | 0.546 | | Marginal $R^2$: 0.468 | | |

**Table S3.** Results of the asymptotic test for the equality of coefficients of variation between colour traits of birds from different habitat types: city centre, residential area, urban part, river corridor and forest. Sample size: great tit n = 201, blue tit n = 108.

| species | colour metric | test statistic | p-value |
|---|---|---|---|
| Great tit | Breast carotenoid chroma | 39.76 | <0.001 |
| | Breast brightness | 8.57 | 0.073 |
| | Wing brightness* | 4.35 | 0.361 |
| | Tail brightness* | 6.21 | 0.184 |
| | Tie area* | 4.29 | 0.368 |
| Blue tit | Breast carotenoid chroma | 1.66 | 0.799 |
| | Breast brightness | 2.32 | 0.677 |
| | Wing inner vane brightness | 0.07 | 0.999 |
| | Wing inner vane UV chroma | 2.15 | 0.709 |
| | Tail brightness | 3.63 | 0.459 |
| | Tail UV chroma | 2.59 | 0.629 |

*Tie area n = 126, wing and tail feather n = 198

**Table S4.** Linear mixed models of great tit and blue tit feather colour traits in relation to tree cover. Significant values are highlighted in bold.

| **GREAT TIT** | **Estimate** | **SE** | **df** | **t value** | **p** | |
|---|---|---|---|---|---|---|
| *Breast carotenoid chroma^2* | | | | | | |
| Intercept | -0.63 | 0.20 | 26.13 | -3.149 | 0.004 | ** |
| **Tree cover** | **0.01** | **0.00** | **194.43** | **3.630** | **<0.001** | *** |
| Sex (M) | 0.22 | 0.15 | 194.54 | 1.494 | 0.137 | |
| Age (older) | 0.11 | 0.15 | 194.99 | 0.743 | 0.458 | |
| Condition index | 0.11 | 0.07 | 194.97 | 1.603 | 0.111 | |
| Random effects: | Variance | | | | | |
| Location | 0.09 | | Conditional $R^2$: 0.194 | | | |
| Residual | 0.87 | | Marginal $R^2$: 0.107 | | | |
| *Breast brightness* | | | | | | |
| Intercept | -0.58 | 0.16 | 14.86 | -3.69 | 0.002 | ** |
| Tree cover | 0.00 | 0.00 | 145.63 | 1.27 | 0.207 | |
| **Sex (M)** | **0.64** | **0.15** | **190.51** | **4.30** | **<0.001** | *** |
| Age (older) | 0.05 | 0.14 | 171.94 | 0.33 | 0.743 | |
| Condition index | 0.06 | 0.07 | 186.46 | 0.90 | 0.371 | |
| Random effects: | Variance | | | | | |
| Location | 0.01 | | Conditional $R^2$: 0.150 | | | |
| Residual | 0.88 | | Marginal $R^2$: 0.136 | | | |
| *Wing brightness* | | | | | | |
| Intercept | 0.37 | 0.18 | 26.05 | 1.99 | 0.057 | |
| Tree cover | 0.00 | 0.00 | 188.05 | -0.69 | 0.490 | |
| **Sex (M)** | **-0.52** | **0.15** | **194.00** | **-3.50** | **<0.001** | *** |
| Age (older) | -0.07 | 0.15 | 193.18 | -0.50 | 0.620 | |
| Condition index | 0.02 | 0.07 | 192.89 | 0.30 | 0.764 | |
| Random effects: | Variance | | | | | |
| Location | 0.06 | | Conditional $R^2$: 0.131 | | | |
| Residual | 0.87 | | Marginal $R^2$: 0.084 | | | |
| *Tail brightness* | | | | | | |
| Intercept | 0.59 | 0.17 | 19.48 | 3.58 | 0.002 | ** |
| Tree cover | 0.00 | 0.00 | 168.50 | 0.02 | 0.986 | |
| **Sex (M)** | **-0.69** | **0.14** | **190.90** | **-4.77** | **<0.001** | *** |
| **Age (older)** | **-0.31** | **0.14** | **180.70** | **-2.15** | **0.033** | * |
| Condition index | 0.12 | 0.07 | 187.40 | 1.76 | 0.080 | . |
| Random effects: | Variance | | | | | |
| Location | 0.03 | | Conditional $R^2$: 0.181 | | | |
| Residual | 0.84 | | Marginal $R^2$: 0.155 | | | |

| | | | | | | |
|---|---|---|---|---|---|---|
| *log Tie area* | | | | | | |
| Intercept | -0.40 | 0.19 | 24.16 | -2.09 | 0.047 | * |
| Tree cover | 0.00 | 0.00 | 118.18 | 1.25 | 0.214 | |
| **Age (older)** | **0.41** | **0.18** | **103.44** | **2.32** | **0.022** | * |
| **Condition index** | **0.23** | **0.09** | **115.15** | **2.52** | **0.013** | * |
| Random effects: | Variance | | | | | |
| Location (Intercept) | 0.01 | | Conditional $R^2$: 0.097 | | | |
| Residual | 0.92 | | Marginal $R^2$: 0.086 | | | |

| **BLUE TIT** | **Estimate** | **SE** | **df** | **t value** | **p** | |
|---|---|---|---|---|---|---|
| *Breast carotenoid chroma* | | | | | | |
| Intercept | -0.48 | 0.23 | 17.02 | -2.05 | 0.056 | . |
| Tree cover | 0.00 | 0.00 | 84.23 | 0.54 | 0.592 | |
| Sex (M) | 0.35 | 0.21 | 100.80 | 1.67 | 0.099 | . |
| Age (older) | 0.32 | 0.20 | 96.99 | 1.62 | 0.109 | |
| Condition index | -0.20 | 0.21 | 91.87 | -0.96 | 0.340 | |
| Random effects: | Variance | | | | | |
| Location | 0.01 | | Conditional $R^2$: 0.065 | | | |
| Residual | 0.98 | | Marginal $R^2$: 0.058 | | | |
| *Breast brightness* | | | | | | |
| Intercept | -0.41 | 0.27 | 38.98 | -1.54 | 0.132 | |
| Tree cover | 0.00 | 0.00 | 100.92 | 0.47 | 0.638 | |
| **Sex (M)** | **0.51** | **0.20** | **98.59** | **2.49** | **0.014** | * |
| Age (older) | -0.17 | 0.19 | 100.57 | -0.89 | 0.377 | |
| Condition index | 0.02 | 0.21 | 100.77 | 0.11 | 0.909 | |
| Random effects: | Variance | | | | | |
| Location | 0.12 | | Conditional $R^2$: 0.178 | | | |
| Residual | 0.89 | | Marginal $R^2$: 0.069 | | | |
| *Wing inner vane brightness* | | | | | | |
| Intercept | 0.22 | 0.28 | 44.51 | 0.78 | 0.442 | |
| Tree cover | 0.00 | 0.00 | 94.66 | 0.24 | 0.815 | |
| Sex (M) | -0.45 | 0.29 | 99.16 | -1.53 | 0.130 | |
| Age (older) | -0.52 | 0.33 | 99.04 | -1.60 | 0.112 | |
| Condition index | 0.03 | 0.21 | 97.83 | 0.16 | 0.873 | |
| **Sex (M): Age (older)** | **0.99** | **0.41** | **96.59** | **2.44** | **0.017** | * |
| Random effects: | Variance | | | | | |
| Location | 0.03 | | Conditional $R^2$: 0.089 | | | |
| Residual | 0.98 | | Marginal $R^2$: 0.059 | | | |

| | | | | | | |
|---|---|---|---|---|---|---|
| *Wing inner vane UV chroma* | | | | | | |
| Intercept | -0.93 | 0.29 | 59.41 | -3.19 | 0.002 | ** |
| Tree cover | 0.01 | 0.01 | 97.13 | 1.66 | 0.101 | |
| **Sex (M)** | **0.79** | **0.33** | **96.33** | **2.37** | **0.020** | * |
| **Age (older)** | **0.88** | **0.17** | **99.88** | **5.08** | **<0.001** | *** |
| Condition index | 0.24 | 0.19 | 100.00 | 1.28 | 0.204 | |
| **Tree cover: Sex (M)** | **-0.01** | **0.01** | **95.88** | **-2.14** | **0.035** | * |
| Random effects: | Variance | | | | | |
| Location | 0.07 | | Conditional $R^2$: 0.299 | | | |
| Residual | 0.76 | | Marginal $R^2$: 0.231 | | | |
| *Tail brightness* | | | | | | |
| Intercept | 0.78 | 0.31 | 69.07 | 2.49 | 0.015 | * |
| Tree cover | 0.00 | 0.01 | 96.58 | 0.43 | 0.669 | |
| Sex (M) | -0.12 | 0.37 | 96.15 | -0.31 | 0.755 | |
| **Age (older)** | **-1.33** | **0.29** | **96.56** | **-4.65** | **<0.001** | *** |
| Condition index | -0.04 | 0.19 | 98.97 | -0.18 | 0.854 | |
| **Tree cover: Sex (M)** | **-0.01** | **0.01** | **95.51** | **-2.03** | **0.045** | * |
| **Sex (M): Age (older)** | **1.05** | **0.36** | **94.61** | **2.93** | **0.004** | ** |
| Random effects: | Variance | | | | | |
| Location | 0.07 | | Conditional $R^2$: 0.303 | | | |
| Residual | 0.74 | | Marginal $R^2$: 0.241 | | | |
| *Tail UV chroma* | | | | | | |
| Intercept | -0.59 | 0.21 | 47.29 | -2.77 | 0.008 | ** |
| **Tree cover** | **-0.01** | **0.00** | **98.41** | **-3.16** | **0.002** | ** |
| **Sex (M)** | **0.74** | **0.28** | **97.58** | **2.68** | **0.009** | * |
| **Age (older)** | **0.77** | **0.15** | **99.27** | **5.04** | **0.000** | *** |
| Condition index | -0.11 | 0.16 | 98.72 | -0.73 | 0.467 | |
| **Tree cover: Age (older)** | **0.01** | **0.005** | **99.27** | **2.15** | **0.034** | * |
| Random effects: | Variance | | | | | |
| Location | 0.02 | | Conditional $R^2$: 0.508 | | | |
| Residual | 0.52 | | Marginal $R^2$: 0.489 | | | |

**Table S5.** Summary results of linear mixed models analysing expression of great tit and blue tit colour traits in relation to impervious surface area cover (%).

| GREAT TIT | Estimate | SE | df | t value | p | |
|---|---|---|---|---|---|---|
| *Breast carotenoid chroma^2* | | | | | | |
| Intercept | -0.15 | 0.17 | 21.15 | -0.85 | 0.408 | |
| **ISA** | **-0.01** | **0.00** | **193.76** | **-2.58** | **0.011** | * |
| **Sex (M)** | **0.30** | **0.15** | **194.99** | **1.98** | **0.049** | * |
| Age (older) | 0.13 | 0.15 | 194.63 | 0.87 | 0.386 | |
| Condition index | 0.13 | 0.07 | 194.40 | 1.76 | 0.080 | . |
| Random effects: | Variance | | | | | |
| Location | 0.07 | | Conditional $R^2$: 0.136 | | | |
| Residual | 0.90 | | Marginal $R^2$: 0.073 | | | |
| *Breast brightness* | | | | | | |
| Intercept | -0.37 | 0.13 | 14.82 | -2.76 | 0.015 | * |
| ISA | -0.01 | 0.00 | 194.99 | -1.67 | 0.096 | . |
| **Sex (M)** | **0.66** | **0.15** | **176.74** | **4.53** | **0.000** | *** |
| Age (older) | 0.06 | 0.14 | 152.80 | 0.44 | 0.661 | |
| Condition index | 0.07 | 0.07 | 185.90 | 1.01 | 0.314 | |
| Random effects: | Variance | | | | | |
| Location | 0.01 | | Conditional $R^2$: 0.130 | | | |
| Residual | 0.89 | | Marginal $R^2$: 0.125 | | | |
| *Wing brightness* | | | | | | |
| Intercept | 0.23 | 0.17 | 21.80 | 1.40 | 0.177 | |
| ISA | 0.00 | 0.00 | 192.91 | 1.44 | 0.152 | |
| **Sex (M)** | **-0.54** | **0.15** | **193.93** | **-3.66** | **0.000** | *** |
| Age (older) | -0.08 | 0.15 | 193.48 | -0.58 | 0.566 | |
| Condition index | 0.02 | 0.07 | 193.23 | 0.23 | 0.816 | |
| Random effects: | Variance | | | | | |
| Location | 0.06 | | Conditional $R^2$: 0.139 | | | |
| Residual | 0.86 | | Marginal $R^2$: 0.080 | | | |
| *Tail brightness* | | | | | | |
| Intercept | 0.55 | 0.15 | 18.94 | 3.80 | 0.001 | ** |
| ISA | 0.00 | 0.00 | 191.93 | 0.75 | 0.455 | |
| **Sex (M)** | **-0.69** | **0.14** | **188.21** | **-4.83** | **0.000** | *** |
| Age (older) | -0.31 | 0.14 | 178.14 | -2.19 | 0.030 | * |
| Condition index | 0.12 | 0.07 | 187.27 | 1.72 | 0.087 | . |
| Random effects: | Variance | | | | | |
| Location | 0.02 | | Conditional $R^2$: 0.180 | | | |
| Residual | 0.84 | | Marginal $R^2$: 0.157 | | | |

| | | | | | | |
|---|---|---|---|---|---|---|
| *Tie area** | | | | | | |
| Intercept | -0.17 | 0.13 | 121.00 | -1.32 | 0.19 | |
| ISA | 0.00 | 0.00 | 121.00 | -0.84 | 0.41 | |
| Age (older) | 0.42 | 0.18 | 121.00 | 2.38 | 0.02 | |
| Condition index | 0.23 | 0.09 | 121.00 | 2.54 | 0.01 | |
| Random effects: | Variance | | | | | |
| Location (Intercept) | 0.00 | | Conditional $R^2$: 0.080 | | | |
| Residual | 0.93 | | Marginal $R^2$: 0.080 | | | |
| **BLUE TIT** | | | | | | |
| *Breast carotenoid chroma* | | | | | | |
| Intercept | -0.30 | 0.22 | 19.32 | -1.38 | 0.185 | |
| ISA | -0.01 | 0.00 | 99.11 | -1.08 | 0.285 | |
| Sex (M) | 0.34 | 0.21 | 97.78 | 1.66 | 0.101 | |
| Age (older) | 0.31 | 0.19 | 95.71 | 1.59 | 0.116 | |
| Condition index | -0.19 | 0.21 | 89.95 | -0.91 | 0.363 | |
| Random effects: | Variance | | | | | |
| Location | 0.00 | | Conditional $R^2$: 0.071 | | | |
| Residual | 0.98 | | Marginal $R^2$: 0.066 | | | |
| *Breast brightness^2* | | | | | | |
| Intercept | -0.15 | 0.25 | 34.58 | -0.61 | 0.543 | |
| ISA | -0.01 | 0.00 | 100.54 | -1.83 | 0.070 | . |
| **Sex (M)** | **0.46** | **0.20** | **98.78** | **2.33** | **0.022** | * |
| Age (older) | -0.19 | 0.19 | 100.50 | -1.02 | 0.310 | |
| Condition index | 0.06 | 0.21 | 100.67 | 0.28 | 0.779 | |
| Random effects: | Variance | | | | | |
| Location | 0.12 | | Conditional $R^2$: 0.205 | | | |
| Residual | 0.86 | | Marginal $R^2$: 0.091 | | | |
| *Wing inner vane brightness* | | | | | | |
| Intercept | 0.22 | 0.26 | 49.91 | 0.84 | 0.408 | |
| ISA | 0.00 | 0.00 | 99.62 | 0.39 | 0.698 | |
| Sex (M) | -0.44 | 0.29 | 99.27 | -1.52 | 0.133 | |
| Age (older) | -0.52 | 0.32 | 99.19 | -1.61 | 0.110 | |
| Condition index | 0.03 | 0.21 | 97.91 | 0.14 | 0.889 | |
| **Sex (M): Age (older)** | **1.01** | **0.41** | **96.90** | **2.49** | **0.015** | * |
| Random effects: | Variance | | | | | |
| Location | 0.03 | | Conditional $R^2$: 0.092 | | | |
| Residual | 0.97 | | Marginal $R^2$: 0.061 | | | |

| | | | | | | |
|---|---|---|---|---|---|---|
| *Wing inner vane UV chroma* | | | | | | |
| Intercept | -0.48 | 0.21 | 33.08 | -2.25 | 0.032 | * |
| ISA | 0.00 | 0.00 | 101.00 | -0.90 | 0.370 | |
| Sex (M) | 0.16 | 0.18 | 100.01 | 0.85 | 0.396 | |
| **Age (older)** | **0.85** | **0.17** | **100.97** | **4.86** | **<0.001** | *** |
| Condition index | 0.22 | 0.19 | 100.75 | 1.16 | 0.248 | |
| Random effects: | Variance | | | | | |
| Location | 0.05 | | Conditional $R^2$: 0.258 | | | |
| Residual | 0.76 | | Marginal $R^2$: 0.210 | | | |
| *Tail brightness* | | | | | | |
| Intercept | 0.73 | 0.25 | 47.92 | 2.92 | 0.005 | ** |
| ISA | 0.01 | 0.00 | 99.95 | 1.45 | 0.150 | |
| **Sex (M)** | **-0.70** | **0.26** | **97.39** | **-2.68** | **0.009** | ** |
| **Age (older)** | **-1.34** | **0.29** | **97.93** | **-4.55** | **<0.001** | *** |
| Condition index | -0.10 | 0.20 | 100.00 | -0.51 | 0.609 | |
| **Sex (M): Age (older)** | **1.01** | **0.37** | **96.11** | **2.76** | **0.007** | ** |
| Random effects: | Variance | | | | | |
| Location | 0.07 | | Conditional $R^2$: 0.266 | | | |
| Residual | 0.78 | | Marginal $R^2$: 0.196 | | | |
| *Tail UV chroma* | | | | | | |
| Intercept | -1.27 | 0.19 | 38.97 | -6.78 | <0.001 | *** |
| **ISA** | **0.01** | **0.00** | **99.35** | **2.34** | **0.022** | * |
| **Age (older)** | **1.48** | **0.18** | **98.26** | **8.21** | **<0.001** | *** |
| **Sex (M)** | **0.71** | **0.15** | **98.61** | **4.61** | **<0.001** | *** |
| condition | -0.14 | 0.16 | 99.99 | -0.88 | 0.380 | |
| ISA: Age (older) | -0.01 | 0.01 | 96.03 | -2.15 | 0.034 | * |
| Random effects: | Variance | | | | | |
| Location | 0.04 | | Conditional $R^2$: 0.515 | | | |
| Residual | 0.52 | | Marginal $R^2$: 0.475 | | | |

*The model with great tit tie area is on the border of singular fit.

### Figures

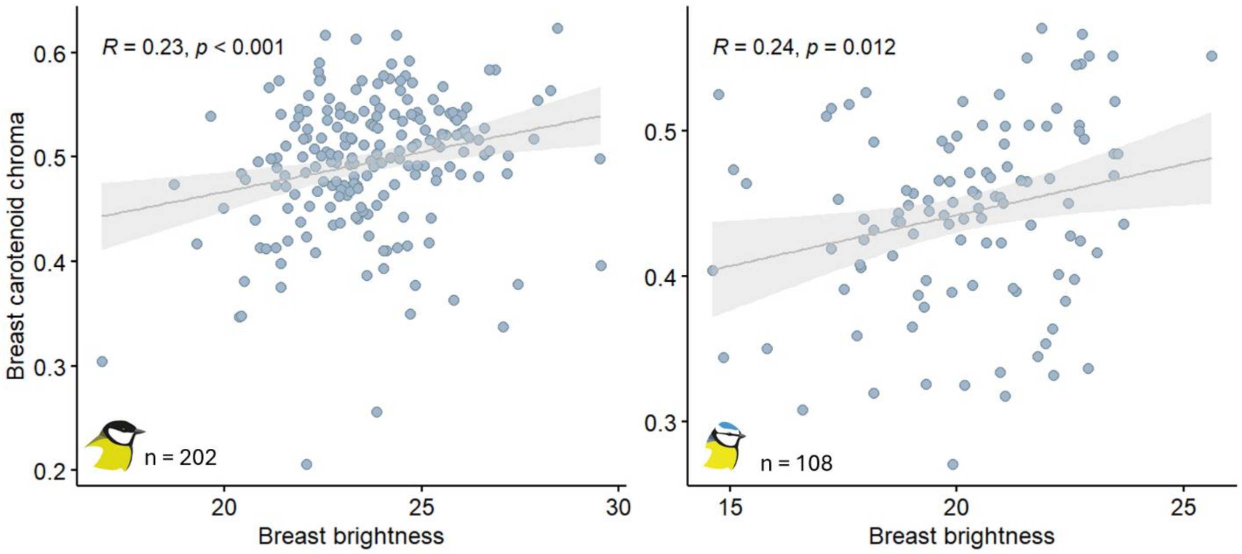

**Figure S1.** Correlations between breast feathers brightness and carotenoid chroma in the great tit (A) and the blue tit (B).

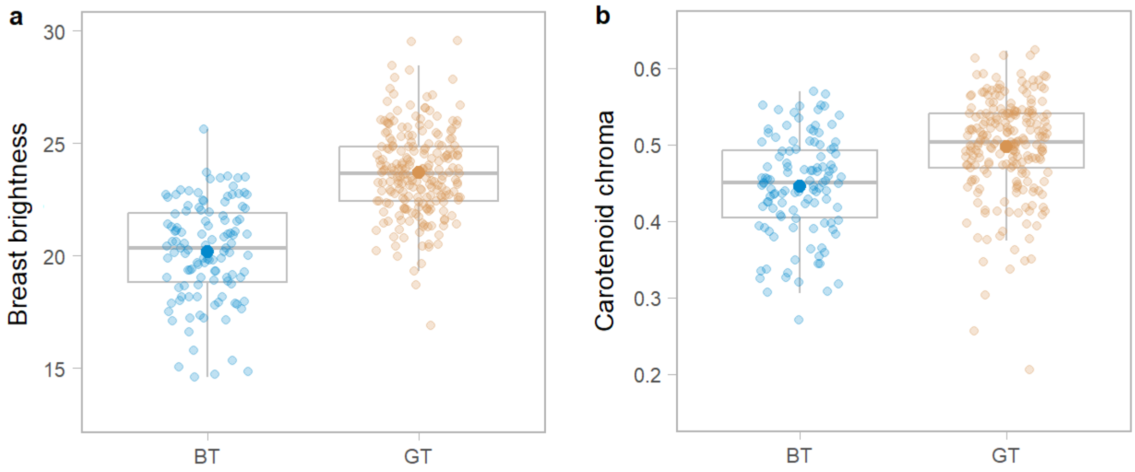

**Figure S2.** Differences in carotenoid-based breast plumage brightness (a) and carotenoid chroma (b) between blue tits and great tits. Sample size: great tit n = 201, blue tit n = 108.

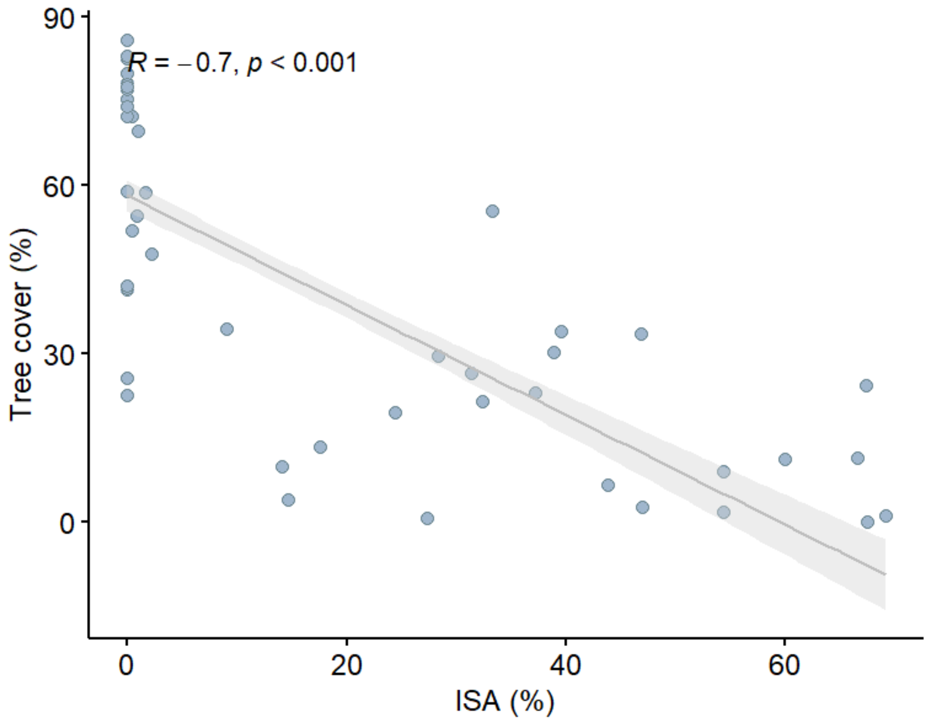

**Figure S3.** Negative relationship between tree cover (%) and impervious surface areas (%) in sampling locations (n = 44).

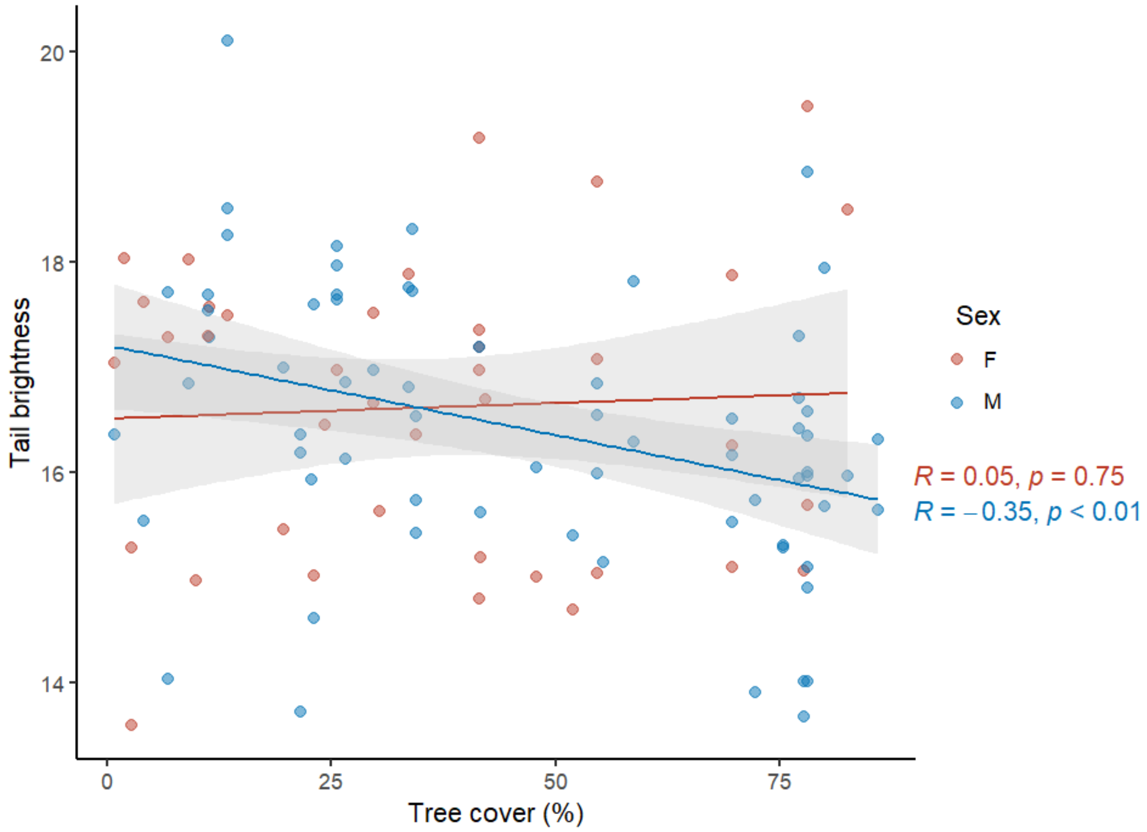

**Figure S4.** Sex specific relationship between blue tit (n = 108) tail brightness and tree cover (%).

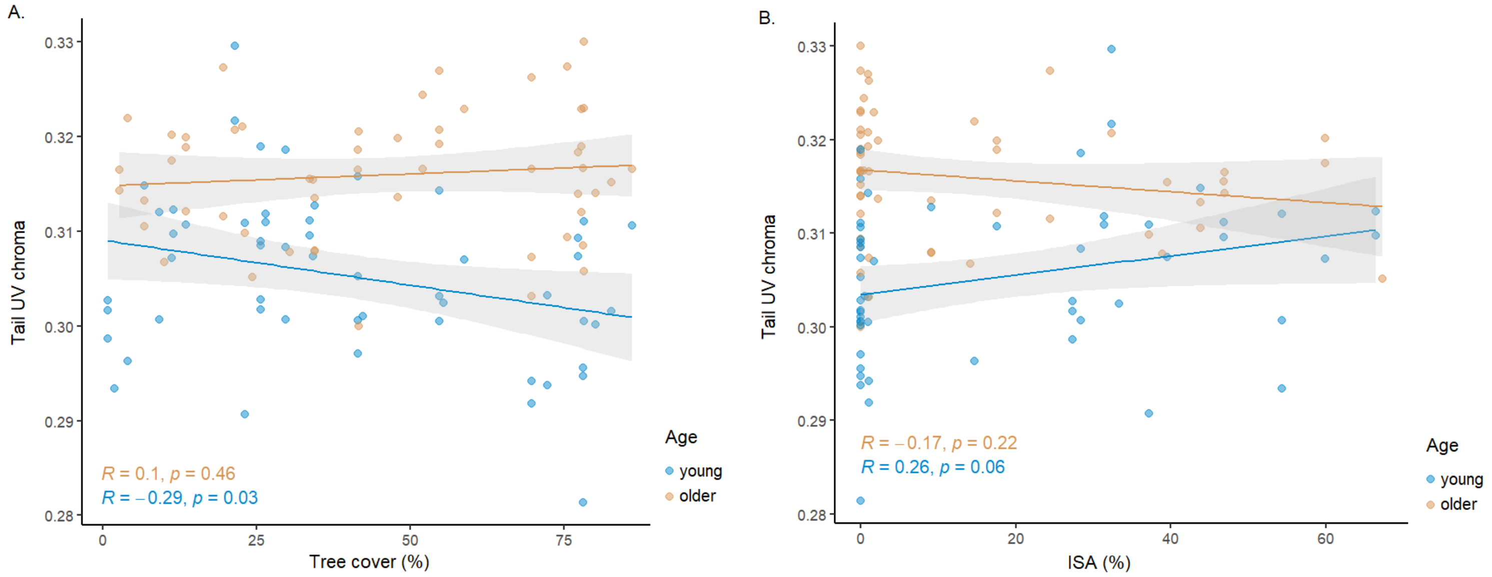

**Figure S5.** Age-specific relationship between blue tit tail UV chroma (n = 108) and tree cover (A.) and ISA (B.).

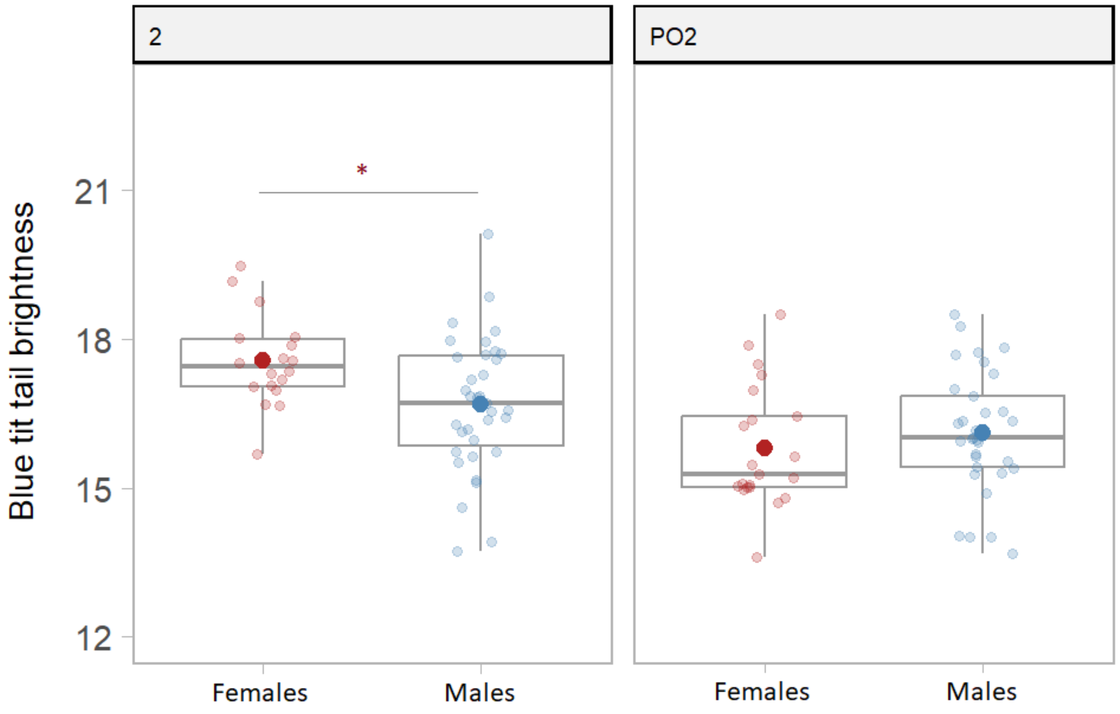

**Figure S6**. Sex differences in blue tit tail brightness. Sample size: birds in their second year :n females = 19, n males = 36; older birds: n females = n 21; n males = 38.

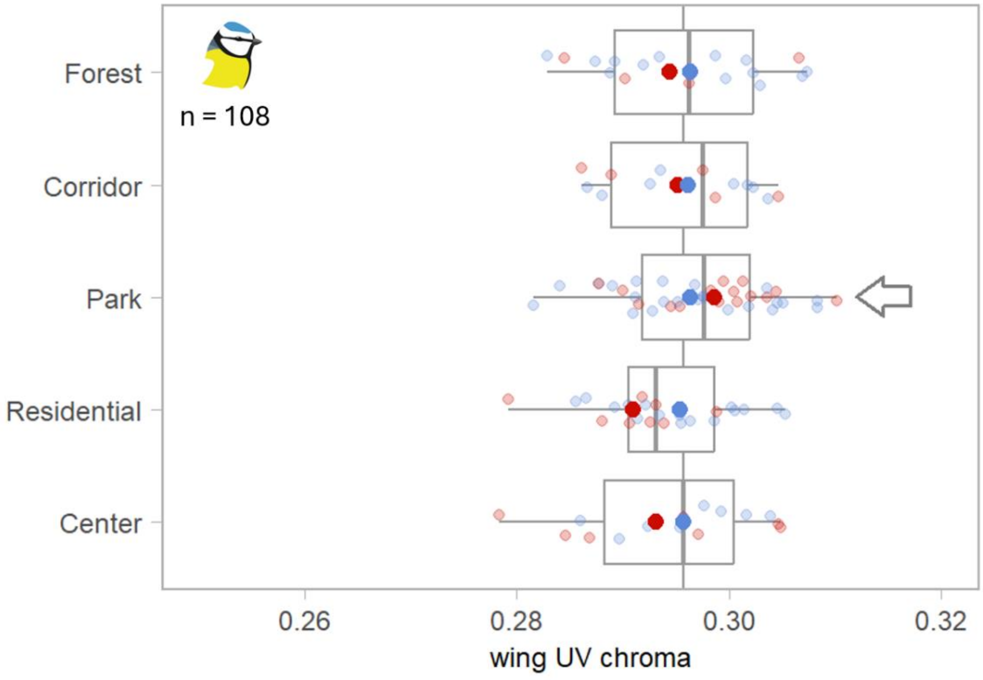

**Figure S7**. Boxplots showing blue tit wing UV chroma in relation to habitat type. Blue and red points represent respectively males and females, horizontal bars indicate data median, bigger dots indicate mean, box edges denote 25%–75% quartiles, whiskers indicate 1.5 IQR. Plotted on raw data.
